# Functional Redundancy in *Candida auris* Cell Surface Adhesins Crucial for Cell-Cell Interaction and Aggregation

**DOI:** 10.1101/2024.03.21.586120

**Authors:** Tristan W. Wang, Dimitrios Sofras, Daniel Montelongo-Jauregui, Telmo O. Paiva, Hans Carolus, Yves F. Dufrêne, Areej A. Alfaifi, Carrie McCracken, Vincent M. Bruno, Patrick Van Dijck, Mary Ann Jabra-Rizk

**Author notes:** Department of Restorative and Prosthetic Dental Sciences, College of Dentistry King Saud bin Abdulaziz University for Health Sciences, Riyadh, Saudi Arabia (Areej A. Alfaifi). Corresponding authors: Mary Ann Jabra-Rizk, Ph.D., University of Maryland-Baltimore, School of Dentistry, 650 W. Baltimore Street, 7 North #7253, Baltimore, Maryland 21201, Patrick Van Dijck, Ph.D., Laboratory of Molecular Cell Biology, Department of Biology, KU Leuven, Leuven, Belgium.

## Abstract

*Candida auris* is an emerging nosocomial fungal pathogen associated with life-threatening invasive disease due to its persistent colonization, high level of transmissibility and multi-drug resistance. Aggregative and non-aggregative growth phenotypes for *C. auris* strains with different biofilm forming abilities, drug susceptibilities and virulence characteristics have been described. Using comprehensive transcriptional analysis we identified key cell surface adhesins that were highly upregulated in the aggregative phenotype during *in vitro* and *in vivo* grown biofilms using a mouse model of catheter infection. Phenotypic and functional evaluations of generated null mutants demonstrated crucial roles for the adhesins Als5 and Scf1 in mediating cell-cell adherence, coaggregation and biofilm formation. While individual mutants were largely non-aggregative, in combination cells were able to co-adhere and aggregate, as directly demonstrated by measuring cell adhesion forces using single-cell atomic force spectroscopy. This co-adherence indicates their role as complementary adhesins, which despite their limited similarity, may function redundantly to promote cell-cell interaction and biofilm formation. Functional diversity of cell wall proteins may be a form of regulation that provides the aggregative phenotype of *C. auris* with flexibility and rapid adaptation to the environment, potentially impacting persistence and virulence.

## INTRODUCTION

The newly emerged nosocomial pathogen *Candida auris* is associated with outbreaks of life-threatening invasive disease worldwide^1–4^. *Candida auris* exhibits several concerning features including persistent colonization of skin and nosocomial surfaces, high transmissibility and unprecedented level of multi-drug resistance^5–11^. In fact, *C*. *auris* is now the first fungal pathogen categorized as an urgent threat by the Center for Disease Control (CDC), making it mandatory to report isolation of *C. auris* in the United States^10,12^. Significantly, the World Health Organization ranks *C. auris* as a critical priority pathogen, highlighting its importance to public health^13^. While virulence factors associated with *C. auris* infections are not fully understood, the fungus shares key characteristics common to *Candida* species including thermotolerance and biofilm formation, although some characteristics are strain-dependent^10,14–16^. Biofilm formation contributes to antifungal tolerance among *Candida* species as a result of drug sequestration, and in *C. auris*, biofilm formation was shown to protect *C. auris* from triazoles, polyenes, and echinocandins^17–19^. One unique growth feature reported in some clinical isolates is cell aggregation, which *in vitro* was associated with differences in drug susceptibility and transcriptional changes induced by exposure to antifungals^20^. Aggregative isolates were also shown to have higher capacity for biofilm formation than non-aggregative isolates^16,18,20–22^.

Fungal cell wall adhesins are crucial for adherence to surfaces and biofilm formation and have been recognized as major virulence factors in *Candida* species^23^. Adhesins also play a fundamental role in interactions of fungal cells with each other enabling switching from a unicellular lifestyle to a multicellular one^18^. In *Candida*, most notably, cell adhesion involves a family of cell surface Als (agglutinin-like sequence) proteins with amyloid-like clusters that activate cell-cell adhesion under mechanical stress^24,25^. Identified polymorphisms enriched in weakly-aggregating strains of *C. auris* were found to be associated with loss of cell surface proteins; furthermore, amplification of the subtelomeric adhesin gene *ALS4* was associated with enhanced adherence and biofilm formation^26,27^. In addition, cell aggregation was shown to increase at higher growth temperatures, suggesting that aggregation is a complex phenomenon that may be linked to the ability to form extracellular matrix and cell surface amyloids^26,28^.

In this study, we aimed to identify unique transcriptional signatures associated with the aggregative phenotype during biofilm growth. As the *in vivo* and *in vitro* situations may have different functional requirements, comparative RNA sequencing analysis was performed on *C. auris* strains grown *in vitro* and *in vivo* using our mouse model of catheter infection. Analysis of differentially regulated genes identified key cell wall adhesin genes to be significantly upregulated in the aggregative strain, and functional analysis of generated null mutants identified an adhesin important for biofilm formation *in vivo*. As complementary roles for diverse adhesins have been reported in *C. albicans*^29^, we aimed to explore adhesin functional redundancy and binding complementation in *C. auris,* which was demonstrated by measuring cell-cell adhesive forces using single-cell atomic force spectroscopy. Functional diversity of cell wall proteins may be a form of regulation providing the *C. auris* aggregative phenotype with flexibility and rapid adaptation to the environment. Therefore, dissecting this aggregative phenotype is crucial for understanding the biology, evolution and pathogenesis of *C. auris*.

## RESULTS

### Transcriptional analysis identifies key cell wall adhesins to be significantly upregulated in an aggregating *C. auris* strain under both *in vitro* and *in vivo* growth conditions

To understand the molecular mechanisms behind the differences observed in the biofilm forming-ability of the two *C. auris* phenotypes, comprehensive RNA-sequencing analysis was performed on cells from *in vitro* grown biofilms. A total of 76 genes were identified to be differentially expressed (LFC ≥ |1|, FDR < 0.01) between the aggregative AR0382 (B11109) and non-aggregative AR0387 (B8441) strains (Fig. 1A); 47 of the genes were more highly expressed in AR0382, whereas 29 genes were more highly expressed in AR0387. Transcriptional analysis of *in vivo* grown biofilms recovered from catheters implanted in mice identified 259 genes that were differentially expressed (LFC ≥ |1|, FDR < 0.01) between AR0382 and AR0387 (Fig.1B); 206 of the genes were more highly expressed in strain AR0382 whereas 53 genes were more highly expressed in AR0387 (Supplementary Table S1 and S2). 23 genes were commonly more highly expressed in AR0382 *in vitro* and *in vivo* (Fig. 1C) (Table S3); among those genes, 5 encode putative adhesins, including B9J08_001458 and B9J08_004112, which have since been annotated as *SCF1 and ALS5*, respectively. Additionally identified were several homologs of *C. albicans* genes with known roles in adhesion including: B9J08_004109 (*IFF4109*) and B9J08_004100 and B9J08_004451 belonging to the *IFF/HYR1* family of adhesins (Fig.1A, B). Interestingly, gene B9J08_002136, an ortholog of the *C. albicans* transcription factor *WOR2* and key regulator of white-to-opaque switching^30^, was significantly underexpressed in strain AR0382 under both growth conditions (Fig.1A, B). In this study, we focused on the two adhesins, Scf1 and Als5.

**Fig. 1.**
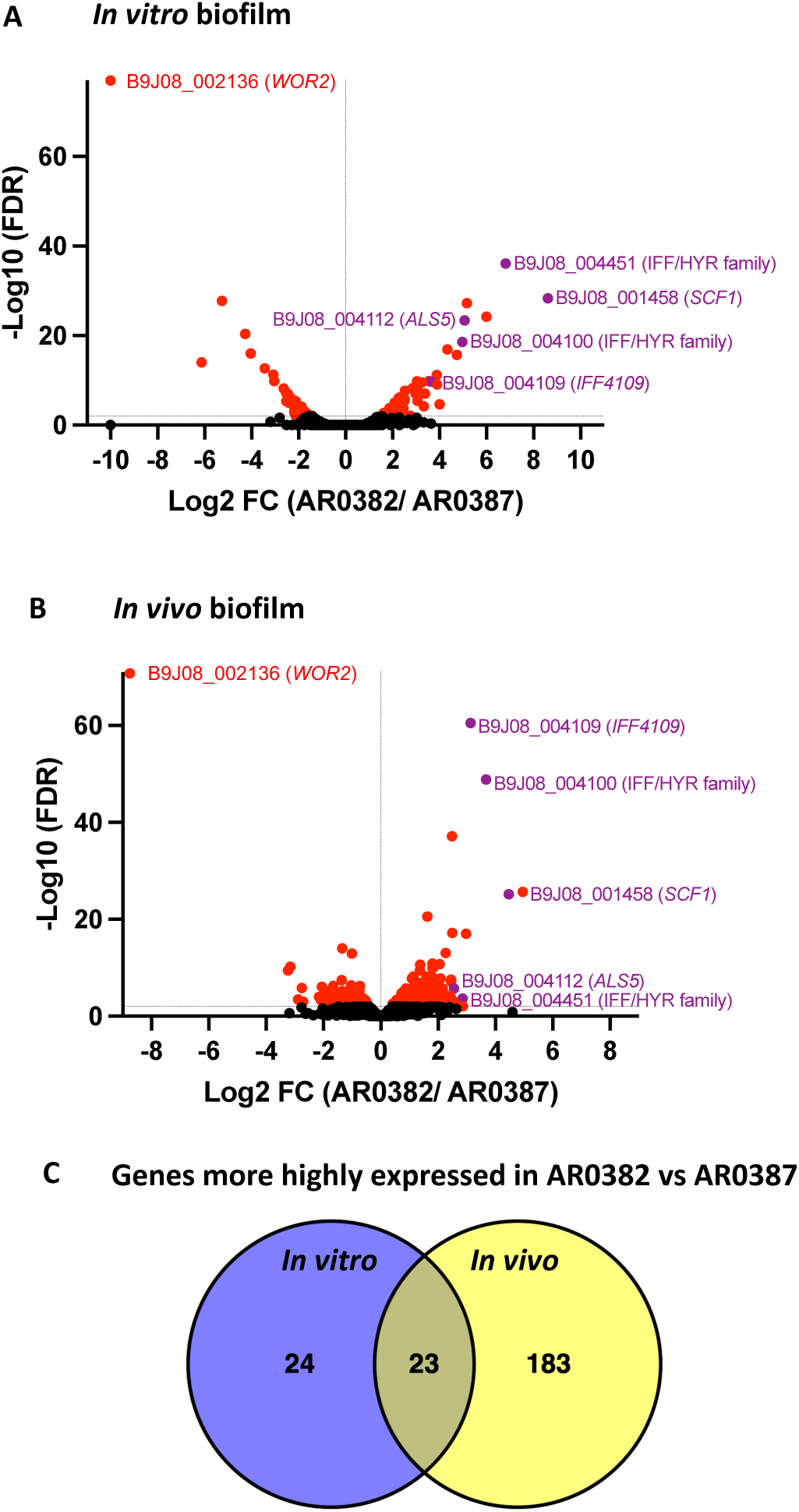
RNA-seq analysis of *in vitro* and *in vivo* grown biofilms depicting genes differentially regulated in the aggregative *C. auris* strain AR0382 compared to the non-aggregative strain AR0387. Volcano plots of comparative differential gene expressions during (A) *in vitro* and (B) *in vivo* biofilm growth. LFC, log (base 2) fold change. FDR, false-discovery rate. *Black*: not statistically significant (FDR > 0.01); *Red*: Statistically significant (FDR < 0.01); *Purple*: Statistically significant and an adhesin. **(C)** Venn diagrams representing the overlap in the numbers of genes that are more highly expressed in strain AR0382 *in vitro* and *in vivo*.

### The *C. auris* Scf1 adhesin contains a Flo11 protein domain and a serine-threonine rich region similar to the *C. albicans* Rbt1 adhesin

We initially identified gene B9J08_001458 as an ortholog of the *C. albicans RBT1* gene in agreement with a previous report^31^; however, this gene has since been renamed *SCF1*^28^. In exploring the similarity between the *C. auris* Scf1 and the *C. albicans* Rbt1 adhesin, comparative analysis of protein domain organization was performed. This revealed a comparable structure for *C. auris* Scf1 to that of the *C. albicans* Rbt1 and the *S. cerevisiae* Flo11 adhesins; specifically, a Flo11 domain and a serine-threonine rich region recognized by Als5, were present in all 3 proteins (Fig. 2).

**Figure 2.**
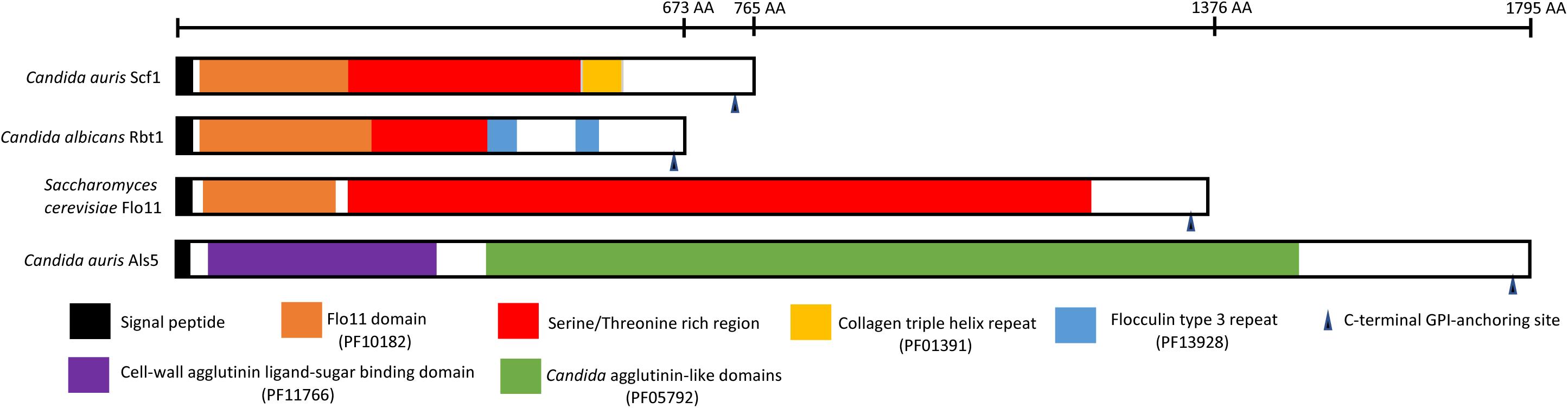
Scf1 adhesin domain organization. Diagram comparing the *C. auris* Scf1 domain structure to that of the *C. albicans* Rbt1 adhesin and the *Saccharomyces cerevisiae* Flo11 depicting a common Flo11 domain and a serine-threonine rich region (>50%) recognized by Als5. Pfam database code is in parentheses; signal peptides and GPI-anchors were predicted using the prediction softwares SignalP 6.0 and NetGPI-1.1, respectively. Functional domains of adhesin proteins were identified *via* InterProtScan (https://www.ebi.ac.uk/interpro/search/sequence/) (accessed February 12, 2024). Uniprot entries: A0A2H1A319 (Scf1); A0A8H6F4R1 (Rbt1); P08640 (Flo11); A0A2H0ZHZ9 (Als5).

### Mutant strain Δ*scf1* but not Δ*als5* is compromised in *in vitro* biofilm formation compared to the wild-type strain

Quantitative evaluation of biofilms based on metabolic activity demonstrated that the Δ*scf1* mutant formed significantly reduced biofilm compared to the wild-type strain. In contrast, the Δ*als5* biofilm was comparable to that of the wild-type (Fig 3A). The non-aggregative AR0387 wild-type strain was severely deficient in biofilm formation compared to AR0382, and deletion of either gene in AR0387 had no additional effect on adhesion and biofilm formation (Fig. S1A).

**Figure 3.**
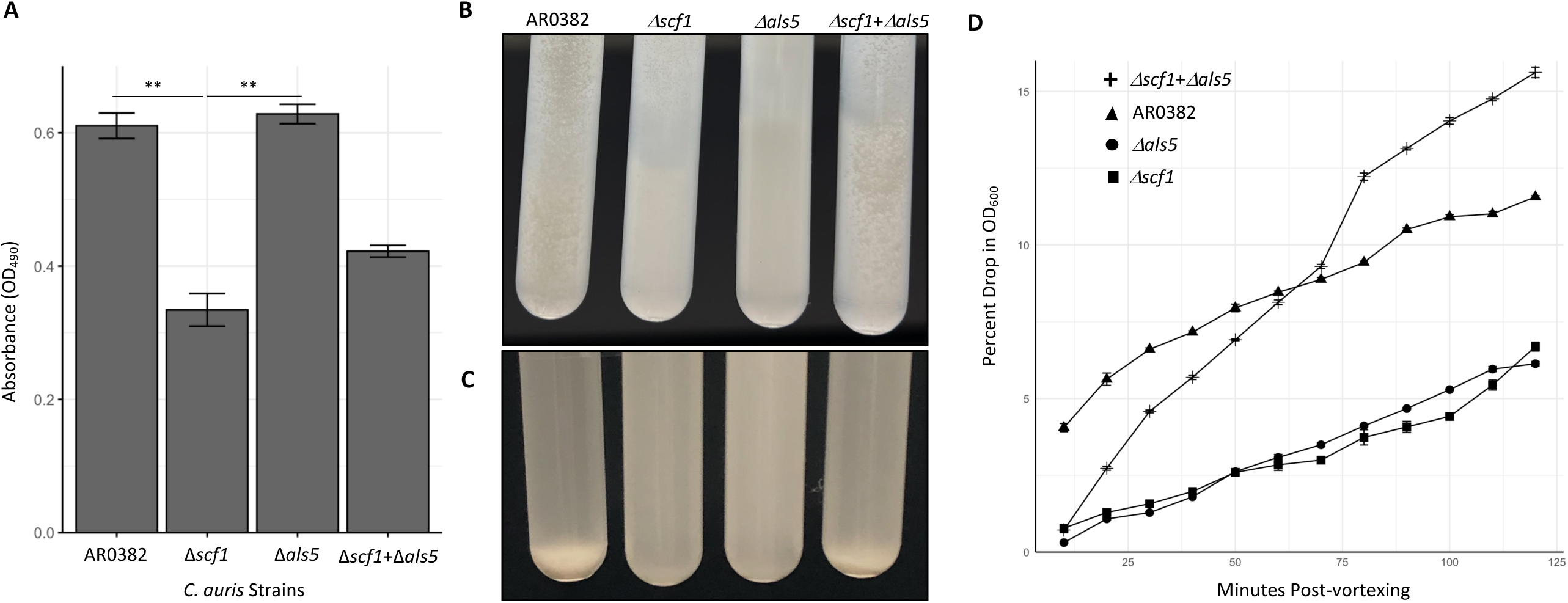
Comparative evaluation of biofilm formation, aggregation and sedimentation rate of Δ*scf1 and* Δ*als5* mutants individually and in combination compared to the wild-type (AR0382). (A) Measurement of the metabolic activity of 24 h biofilms based on values of OD_490_ comparing wild-type AR0382 to Δ*scf1 and* Δ*als5* mutants and Δ*scf1*+Δ*als5* combination. Statistical analysis was performed by one-way ANOVA and post-hoc Tukey test with *p*-values representing significant differences. Bar-plots show mean and SEM of *n* = 3 biological replicates, each as an average of 4 technical replicates. *P* = 2.61×10^-3^, 1.75×10^-3^. **(B)** Cell aggregation 2 min after vigorous vortexing and **(C)** 10 min post-vortexing. **(D)** Measurement of rate of cell sedimentation over 2 hr. Values represented are mean OD plus SEM of three technical replicates. **0.001 < P ≤ 0.01.

### Both mutant strains Δ*scf1* and Δ*als5* are significantly deficient in aggregation

Following vortexting of cell suspensions, the AR0382 wild-type strain formed large aggregates rapidly settling into a sediment (Fig. 3B, C). In contrast, mutants lacking the Scf1 or Als5 adhesins formed no or minimally visible aggregates and cells remained mostly suspended (Fig. 3B, C). Comparative measurement of sedimentation rates of aggregates based on drop in absorbance readings over time demonstrated that unlike with the Δ*scf1* and Δ*als5* mutants, the drop in absorbance for the wild-type strain was dramatic (Fig. 3D). No aggregation was seen with cells of the AR0387 wild-type strain (Fig. S1B-C).

### Scf1 and Als5 adhesins have complementary and redundant roles in cell-cell adherence and aggregation

In order to explore whether the two highly expressed adhesins in the aggregative strain have complementary roles, the two mutant strains were mixed and cell-cell adherence and coaggregation were monitored visually and quantitatively. Where individually both mutants failed to aggregate, in combination, cells co-adhered strongly, forming aggregates comparable to those formed by the wild-type strain (Figs. 3B-D).

### Confocal Laser Scanning Microscopy (CLSM) and Scanning Electron Microscopy (SEM) imaging reveal significant differences in biofilm architecture for Δ*scf1* and Δ*als5* compared to the wild-type and Δ*scf1*+Δ*als5* mixed biofilm

CLSM images revealed significant differences in the extent and structure of biofilms formed by the wild-type and Δ*scf1*; where the wild-type biofilm consisted of dense matrix and cell aggregates, the Δ*scf1* biofilm was patchy and less dense (Fig. 4). Although Δ*als5* formed a substantial biofilm, it was not as dense or aggregative as the wild-type. In contrast, biofilm formed by combination of Δ*als5* and Δ*scf1* was comparable to that of the wild-type. SEM analysis revealed similar biofilm structures where wild-type and Δ*als5+*Δ*scf1* mixed biofilms consisted of piles of cell aggregates, and biofilms of Δ*als5* and Δ*scf1* were homogenous consisting mostly of single layer of cells (Fig. 5).

**Figure. 4.**
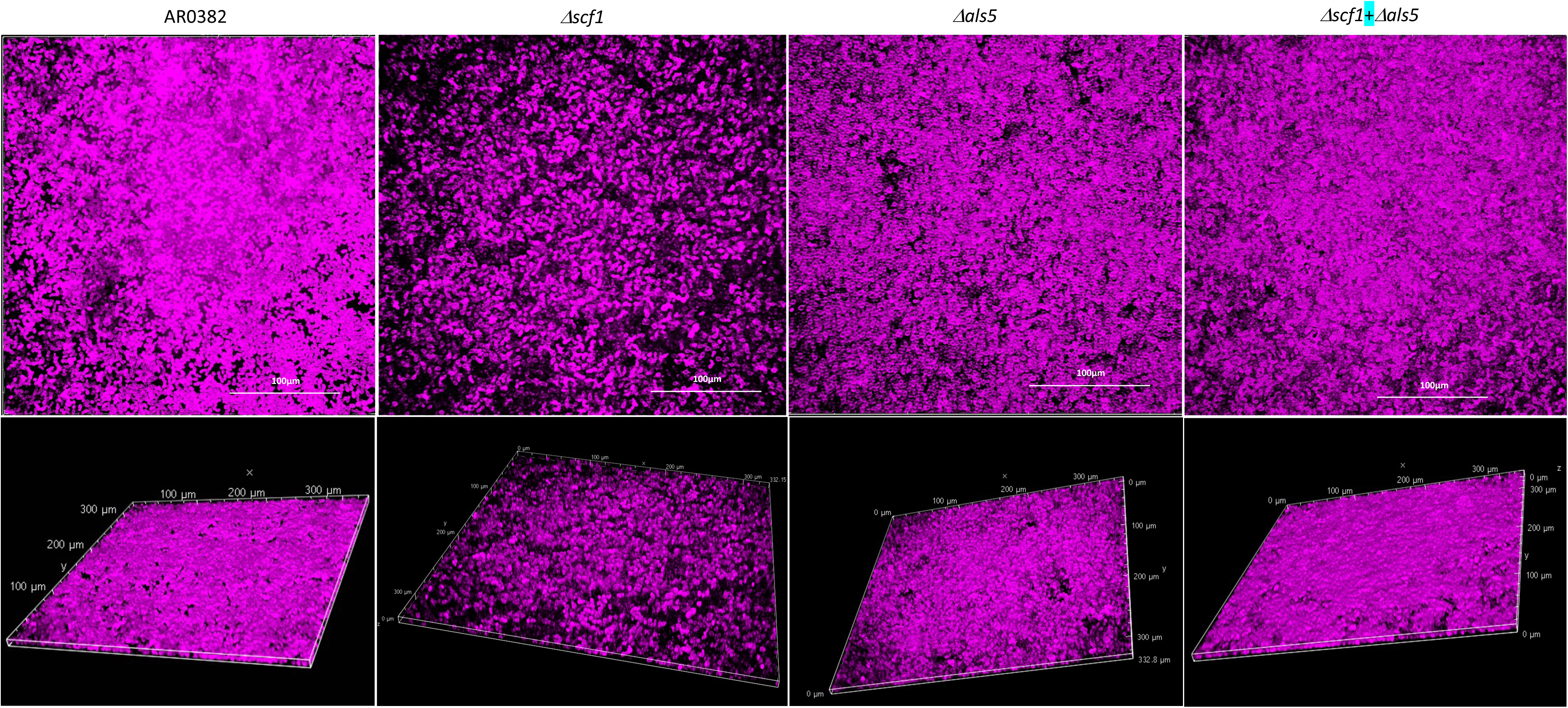
Representative images from confocal laser scanning microscopy analysis of biofilms formed by the *C. auris* AR0382 wild-type (WT) strain and the Δ*scf1* and Δ*als5* mutants grown individually and in combination (Δ*scf1+*Δ*als5)*. Z-stack reconstructions of biofilms stained with polysaccharide stain concanavalin A (fuchsia).

**Figure. 5.**
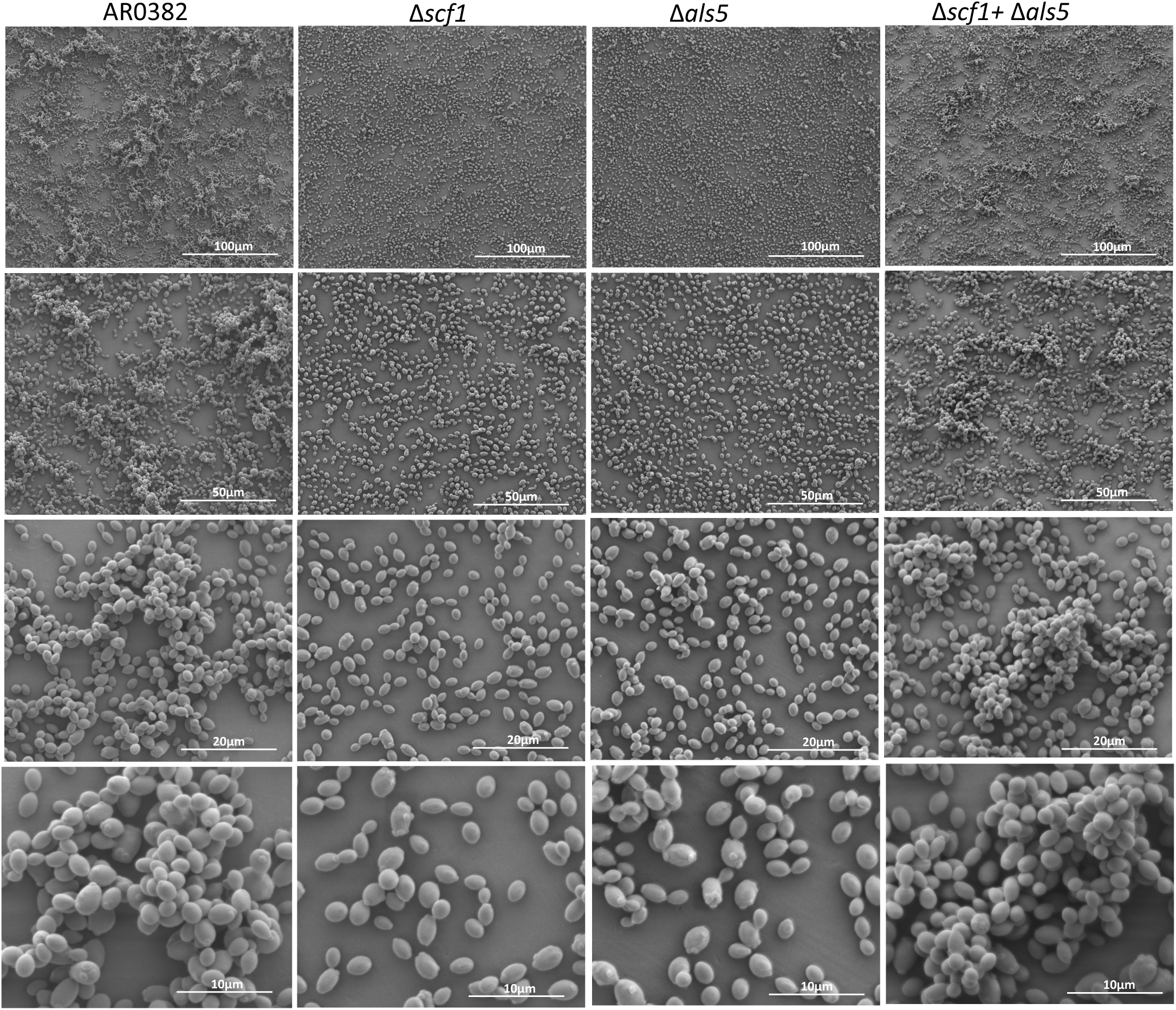
Representative images from scanning electron microscopy analysis. 24 h biofilms formed by the *C. auris* AR0382 wild-type (WT) strain and the *Δscf1* and *Δals5* mutants grown individually and in combination (Δ*scf1*+Δ*als5)*.

### Atomic force microscopy (AFM) reveals major differences in cell-cell adhesion forces between the different strains

Force-distance curves recorded by AFM^32^ between two AR0382 wild-type cells featured a large adhesion force peak averaged at 338 ± 219 pN (mean ± standard deviation (SD), n = 1567 adhesive curves from 6 cell combinations) (Fig. 6A, B). Moreover, some force profiles showed sawtooth patterns with multiple force peaks in the 200-500 pN range, which could be attributed to the sequential unfolding of the tandem repeat domains of Als proteins^33^ (Fig. 2). Interestingly, a wide distribution of adhesion forces, composed of both weak and strong forces was observed for this strain. In the non-aggregative AR0387 strain however, intercellular adhesion was essentially non-existent (4%) and only weak forces of 96 ± 29 pN (305 adhesive curves from 4 cell pairs) were measured (Fig. S1 D-F). In contrast to AR0382, significant decrease in adhesion frequency was observed for the Δ*als5* strain (from 80% to 30%) (Fig. 6C), where force curves featured only weak adhesion forces of 127 ± 27 pN (n = 608 adhesive curves from 6 cell pairs) (Fig. 6B), and sawtooth patterns were not observed (Fig. 6A). Similar intercellular adhesion forces were measured for the Δ*scf1* strain (111 ± 30 pN, n = 731 adhesive curves from 6 cell pairs), and an adhesion frequency slightly higher than what was observed for the Δ*als5* strain (46%) Fig. 6C). Finally, adhesion was also probed between cells of the Δ*als5* and Δ*scf1* mutants; even though a mean adhesion frequency of 54% was registered (Fig. 6C), half of the cell pairs probed exhibited adhesion frequency in the same range of what was observed for the AR0382 wild-type strain. Despite this difference, intercellular adhesion forces of 132 ± 43 pN (n = 1146 adhesive curves from 8 cell combinations) were measured for the Δ*als5+*Δ*scf1* experiment (Fig. 6A, B).

**Figure 6.**
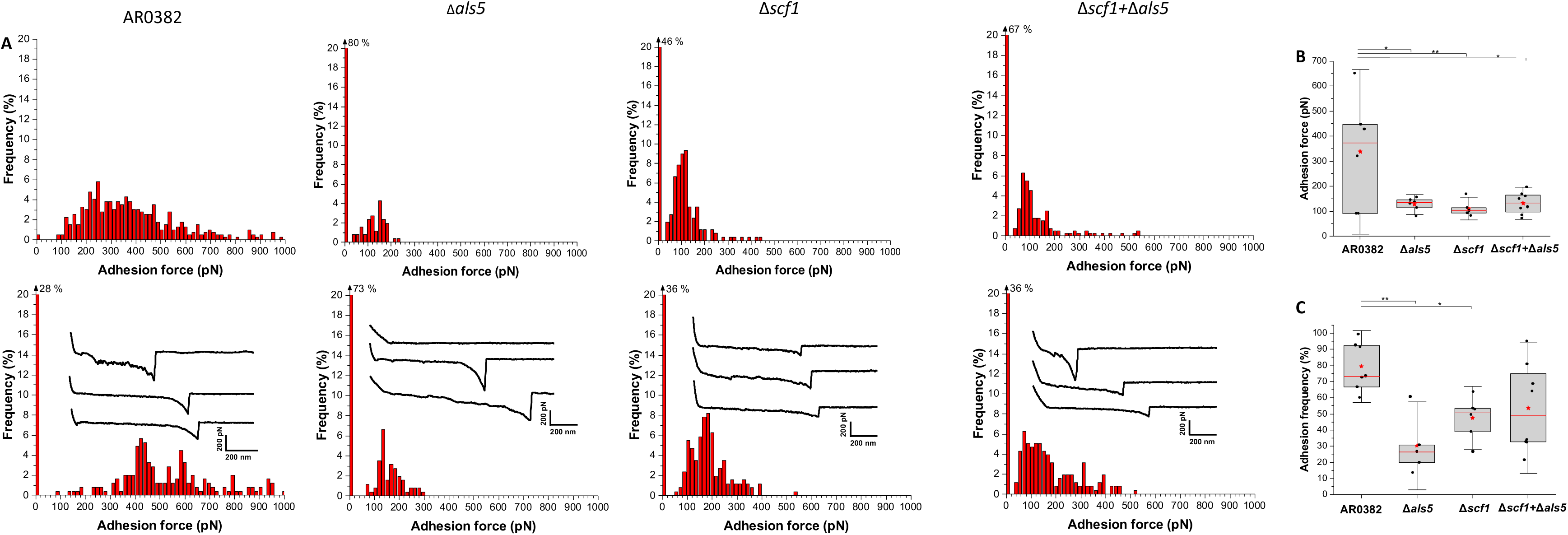
Single-cell force spectroscopy of *C. auris* cell-cell adhesion. (A) Adhesion force histograms with representative retraction profiles (inset) obtained for the interaction between AR0382 wild-type cells, cells of Δ*als5,* cells *of* Δ*scf1* and between cells of Δ*als5* and Δ*scf1 (*Δ*als5*+Δ*scf1);* 2 representative cell pairs are shown for each strain. **(B)** Adhesion force boxplots show data on *n* = 6 pairs of AR0382 cells, Δ*als5* cells, and Δ*scf1* cells and *n* = 8 cell pairs combining Δ*als5* and Δ*scf1*. Statistical analysis was performed by one-way ANOVA and post-hoc Tukey test with *p*-values representing significant differences. *P* = 1.42×10^-2^, 7.69×10^-3^, 1.01×10^-2^. **(C)** As in **(B)**, adhesion frequency boxplots show data on *n* = 7 pairs of AR0382 cells, *n* = 5 pairs of Δ*als5* cells*, n =* 6 pairs of Δ*scf1* cells and *n* = 8 pairs between Δ*als5*+Δ*scf1* cells. *P*=1.71×10^-3^, 3.77×10^-2^. Red stars represent the mean values, red lines are the medians, boxes are the 25−75% quartiles and whiskers the standard deviation from mean. *0.01 < *P* ≤ 0.05, **0.001 < *P* ≤ 0.01.

### SEM analysis of catheters implanted in mice showed impaired *in vivo* biofilm formation in Δ*scf1* mutant compared to wild-type strains

SEM imaging of Infected catheters recovered from mice (Fig. 7A) revealed significant differences in density and architecture of biofilms formed within catheter lumens. The AR0382 wild-type strain formed a robust biofilm consisting of aggregates of cells; in contrast, biofilms within catheters infected with Δ*scf1* were scarce with patches consisting primarily of extracellular matrix with fewer yeast cells in single layers and no or minimum cell aggregates, comparable to that formed by the non-aggregative AR0387 wild-type strain (Fig. 7B).

**Figure. 7.**
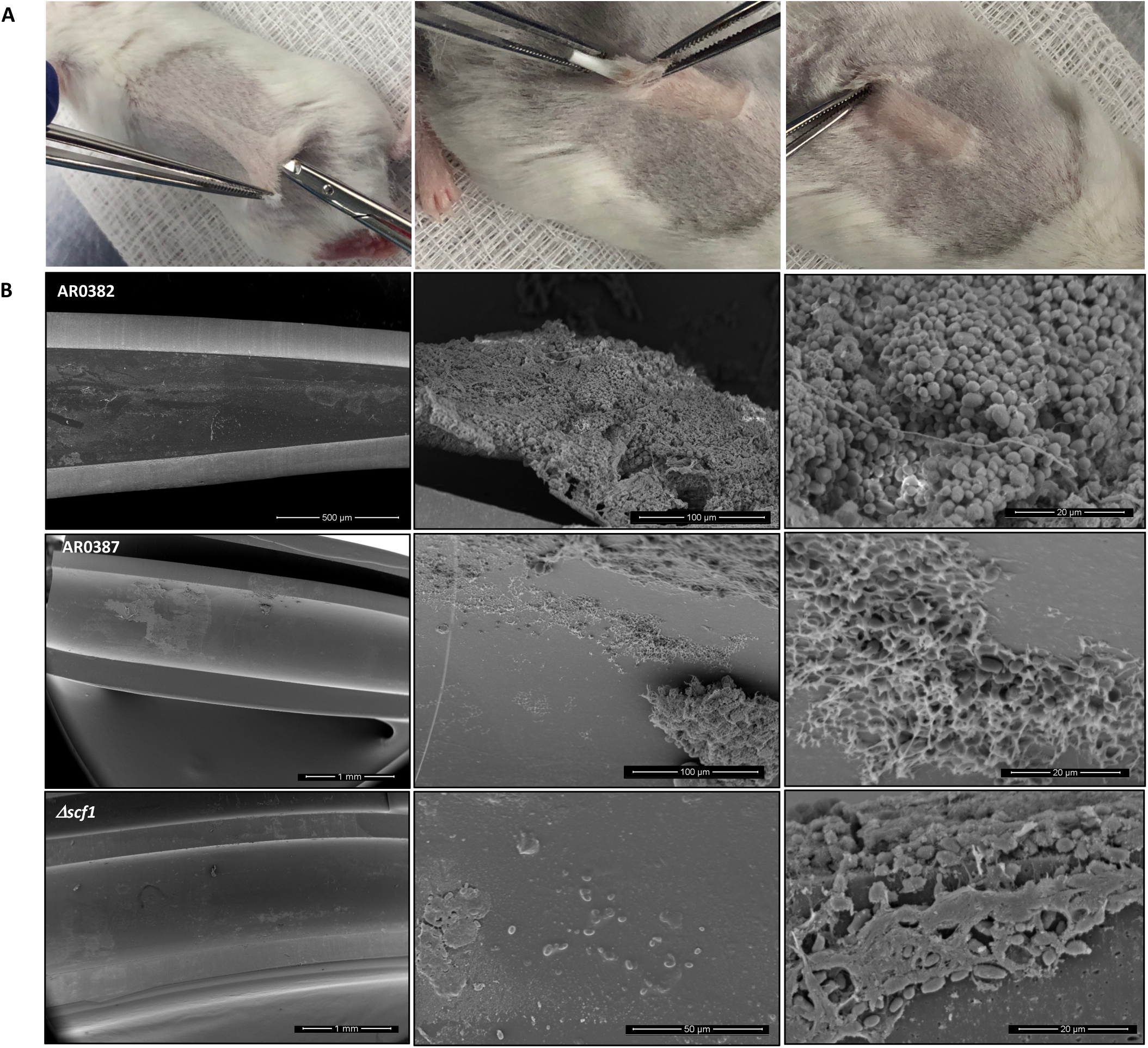
Infection and biofilm formation in catheters implanted in mice. (A) A small incision is made in a shaved area in the dorsum of anesthetized mice and catheter fragments (0.5 cm) are inserted within a formed subcutaneous tunnel. **(B)** Scanning electron microscopy of explanted catheters. Representative low- and high-magnification SEM images demonstrating mature biofilm formed within lumen of catheters infected with AR0382 wild-type strain consisting of aggregates of yeast cells.

## DISCUSSION

*Candida auris* avidly adheres and forms biofilms on indwelling medical devices such as intravascular catheters, an important risk factor for systemic infection. A striking morphological feature of some *C. auris* isolates is their capacity to aggregate and form strong biofilms^18,27^. In this study, we performed comprehensive comparative analysis of biofilms formed by strains exhibiting a high and low aggregation phenotype under *in vitro* and *in vivo* growth conditions. First, our analysis focused on identifying genes that were more highly expressed in the aggregative strain both *in vitro* and *in vivo*, specifically those with predicted roles in adhesion and biofilm formation based on functional homology in other *Candida* species. Most notable among the genes that are more highly expressed in the aggregative strain were B9J08_001458 and B9J08_004112, which encode homologs of *C. albicans RBT1* and *ALS5*, respectively^31,34,35^. Recently, B9J08_001458 was described as unique to *C. auris* and was named *SCF1* by Santana *et al.*^28,36^; protein domain analysis shows that both Scf1 and *Saccharomyces cerevisiae* Flo11p share a N-terminal Flo11 domain^28^. The Flo adhesin family initially discovered in brewer’s yeast *(S. cerevisiae)* has the ability to form cellular aggregates induced by shear force^18,29^. Interestingly, we identified a Flo11 domain in the *C. albicans* Rbt1 in the N-terminal domain and sequence comparisons demonstrated high similarities between the Flo11 domains of the *C. auris* Scf1 and *C. albicans* Rbt1 (Fig. 2). The *C. albicans* Rbt1 adhesin is involved in cell-cell adhesion and overexpression of *RBT1* in *C. albicans* was shown to trigger the clustering of other cell surface proteins harboring aggregate-forming sequences such as Hwp1, by forming intermolecular bonds^31,37^. Further, Rbt1 is related to the Hwp1 and Hwp2 cell wall proteins that play distinct but overlapping roles in *C. albicans* for promoting biofilm formation^38^. In fact, the Hwp1 protein possesses an internal serine-threonine-rich region with a critical role in cell-cell adhesion and biofilm formation^39^. Therefore, we propose that Scf1 functions as an adhesin in a similar manner to the *C. albicans* Rbt1 and Hwp1.

Fungal cellular aggregation is proposed to occur as a result of a global cell surface conformational shift^40^. Therefore, we aimed to investigate the contribution of the *C. auris* Scf1 in relation to other expressed adhesins, primarily the co-upregulated Als5. Heterologous expression of the *C. albicans* Als5 at the surface of *S. cerevisiae* was shown to result in Als5-mediated adhesion followed by formation of multicellular aggregates, which was not observed when *ALS5* was expressed at reduced levels^41,42^. In exploring the mechanism driving Als5-mediated intercellular adhesion, a study described an aggregation mechanism whereby amyloid core sequences in Als proteins trigger the formation of cell surface adhesion nanoclusters, facilitating strong interactions between adhesins on opposing cells^25^. Interestingly, in *C. albicans*, Als5 adhesion was shown to be mediated by recognition of a minimum of four accessible contiguous threonine and serine residues^43,44^. Our analysis of the Scf1 protein sequence identified the presence of five contiguous Als5-recognized threonine-serine rich domains comparable to that in the *C. albicans* Hwp1 and Rbt1, further supporting the functional similarity of Scf1 to this class of *C. albicans* adhesins (Fig. 2). This degenerate “recognition system” among adhesins would provide *C. auris* with a plethora of target proteins for adherence^44^.

Interestingly, complementary roles for *C. albicans* Hwp1 and Als1/3 in biofilm formation have been described by Nobile *et al.*^29^, whereby a mixture of biofilm-defective *HWP1* and *ALS1/3* mutants could form a hybrid biofilm. Hence similarly, despite their sequence divergence, we posit that in *C. auris*, Als5 and Scf1 may function redundantly to promote cell-cell interaction and biofilm formation (Fig. 8). To that end, we generated gene deletion strains of *C. auris SCF1* and *ALS5* in the aggregative strain AR0382. Interestingly, phenotypic evaluations demonstrated reduced adhesion and biofilm formation *in vitro* for the Δ*scf1* mutant, but not for the Δ*als5* mutant compared to the wild-type strain (Fig. 3A). Individually, cells of Δ*scf1* and Δ*als5* lost aggregation capability, but aggregation was restored when combined (Fig. 3B-D). This aggregation was also demonstrated by SEM analysis, revealing a confluent mixed biofilm comprised of heaps of co-adhering cells, comparable to that seen with the wild-type (Fig. 5). Interestingly however, although based on assessment of metabolic activity the Δ*als5* biofilm was comparable to that of the wild-type, SEM biofilm imaging revealed dramatic structural differences. These observations corroborate a previous report that Als5 is not crucial for adherence to abiotic surfaces^28^. However, here we show that this adhesin is necessary for mediating cell-cell adherence. Due to the observed reduction in the ability of Δ*scf1* to form biofilm *in vitro*, we then tested this mutant in our mouse model to evaluate biofilm formation *in vivo*. In contrast to the dense aggregative biofilm formed by the AR0382 strain, SEM imaging of the biofilms within catheters revealed a minimal biofilm formed by Δscf1, comparable to that of the AR0387 strain consisting primarily of single layers of yeast cells, (Fig. 7).

**Figure. 8.**
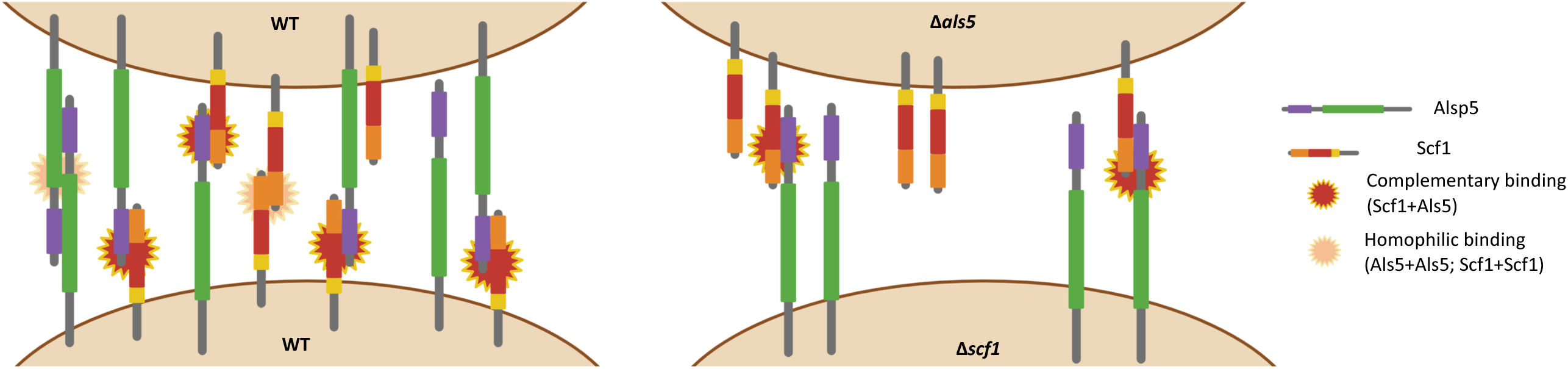
Hypothetical mechanistic model depicting complementary Scf1/Als5 binding. (left) Adherence between wild-type (WT) cells involving Scf1+Als5 complementary binding and homophilic interactions between Als5+Als5 and Scf1+Scf1; (right) Complementary binding between Scf1 and Als5 on the Δ*als5* and Δ*scf1* mutant cells, respectively. Domain designations and colors are consistent with those in Fig. 2.

The strong cell-cell affinities between the Δ*scf1* and Δ*als5* mutants were assessed by measuring adhesion forces using single-cell force spectroscopy (Fig. 6 and Fig. S1). With the wild-type strain, a wide distribution of adhesion forces composed of both weak and strong forces were detected, indicative of a complex binding mechanism that involves a combination of single and multiple molecular bonds. The involvement of the Als5 and Scf1 adhesin in cell-cell adhesion was demonstrated by the significant decrease in adhesion frequency observed between cells of the Δ*als5* and Δ*scf1* strain, which was partially restored by mixing the two deletion strains. High forces were not completely restored however when probing Δ*als5* cells with Δ*scf1* cells (and *vice versa*), indicating that *C. auris* cell-cell adhesion not only involves a combination of single and multiple Als5-Scf1 bonds, but also Als5-Als5 and Scf1-Scf1 homophilic bonds, and potentially other mechanisms (Fig. 8). In fact, it is well-documented for *C. albicans* that Als5 proteins are able to form intercellular amyloid bonds through their T domains to promote biofilm formation^25,33,45,46^. Additionally, since the *C. albicans* Rbt1 was also shown to be capable of forming amyloid bonds^37,47^, it is tempting to speculate that similar homophilic binding might similarly occur with the *C. auris* Scf1 adhesin. Combined, these findings indicate that Als5 and Scf1 undergo a complementary heterophilic binding reaction that supports *C. auris* cell-cell adherence, critical for intraspecies adhesin interactions and promoting formation of monospecies biofilms.

Collectively, our findings demonstrated significant *in vitro* and *in vivo* transcriptional changes associated with the *C. auris* aggregative form impacting cell wall adhesins that although with little similarity, may have complementary roles, and function redundantly to promote cell-cell interaction and biofilm formation (Fig. 8). Functional diversity of cell wall proteins may be a form of regulation providing the *C. auris* aggregative phenotype with flexibility and rapid adaptation to the environment, potentially impacting persistence and virulence

## METHODS

### Strains and growth conditions

The *C. auris* wild-type strains AR0382 (B11109) and AR0387 (B8441) from the CDC AR-panel were used as wild-type strains in this study. We have previously characterized these strains and designated AR0382 as aggregative/high biofilm former and AR0387 as non-aggregative/low biofilm former^16^. Both isolates were confirmed to belong to clade I (East Asian) based on Carolus *et al.*^48^ and were isolated in Pakistan; AR0382 was recovered from a burn wound and AR0387 from blood. Mutant strains of *C. auris* genes B9J08_001458 and B9J08_004112 in the AR0387 and AR0382 backgrounds were generated in this study. These genes have since been named *SCF1* and *ALS5,* respectively^27,28^. Isolates were grown overnight in yeast peptone dextrose broth (YPD) (Difco Laboratories) at 30°C, washed in Phosphate Buffered Saline (PBS) and resuspended in PBS to final cell density needed.

### *In vitro* transcriptional analysis of AR0387 and AR0382 biofilms using RNA-sequencing

Biofilms of both wild-type strains were formed in 6-well plates in RPMI 1640-HEPES media (Invitrogen) at 37°C for 24 h. Following incubation, wells were rinsed with PBS and biofilms were scraped. Recovered cells were snap-frozen on dry ice and ethanol, allowed to thaw at room temperature (RT), then incubated for 30 min at 37°C in “*digestion buffer*” containing 100 U/ml of lyticase and RNAse inhibitor in TRIS-EDTA 1x buffer. RNA was extracted in 1 ml of TRI Reagent™ Solution (Ambion, Invitrogen; Carlsbad, CA) using bead-beating for 30 min at RT followed by purification using Direct-zol RNA Miniprep kit (Zymo Research; Tustin, CA). Eluted RNA was analyzed in a Nanodrop Lite (Thermo Scientific). Total RNA was subjected to rRNA depletion with the Ribominus Eukaryote Kit. All RNA-seq libraries (strand-specific, paired end) were prepared with the TruSeq RNA sample prep kit (Illumina). One hundred nucleotides of sequence were determined from both ends of each cDNA fragment using the Novaseq platform (Illumina). Sequencing reads were aligned to the reference genomes (*C. auris* strain B8441) using HISAT2^49^, and alignment files were used to generate read counts for each gene; statistical analysis of differential gene expression was performed using the DEseq package from Bioconductor^50^. A gene was considered differentially expressed if the absolute log fold change was greater than or equal to 1 and the FDR value for differential expression was below 0.01. The RNA-seq analysis was performed in biological triplicate. Given the limited annotation of the *C. auris* genome, some of the gene names reported are based on homology to *C. albicans* genes. For genes with no recognizable orthologs, the original systematic *C. auris* gene designation is provided.

### *In vivo* transcriptional analysis of AR0387 and AR0382 biofilms formed within catheters implanted in mice using RNA-sequencing

All animal experiments were conducted at the AAALAAC accredited Animal Facility of the University of Maryland, Baltimore and were approved by Animal Care and Use Committee. Three-month-old female Balb/c mice (Jackson Laboratory) were housed at a maximum of 5 per cage, weighed and closely monitored for any signs of distress. A modified model previously described by Kucharíková *et al.*^51^ was used; 0.5 cm fragments of polyurethane triple-lumen central venous catheters (Jorgensen Laboratories) pre-coated overnight with fetal bovine serum (Gibco™) were incubated with 1x10^8^ cells/ml cell suspensions in PBS for 1.5 h at 37°C, rinsed and kept on ice until implanted. For each experimental set, *in vitro*-infected catheters were processed for assessment of microbial recovery. Mice were anesthetized with 0.5 ml intraperitoneal injections of tribromoethanol (TBE) solution (250 mg/kg; Sigma-Aldrich); dorsum of mice was shaved and a small incision made aseptically and a subcutaneous tunnel was created allowing for insertion of up to 5 pieces of pre-inoculated catheters (Fig. 7A). Incisions were sealed using 3M Vetbond™ tissue glue and lidocaine analgesic gel was applied. Biofilms were allowed to form within catheters for 72 h then animals were euthanized by CO_2_ inhalation followed by cervical dislocation. Catheters were collected in RNAlater buffer, aseptically fragmented, sonicated in RNAse free water and cells from all catheters recovered from each mouse were pooled by centrifugation. RNA-sequencing was performed as described above. The AR0382 group contained three biological replicates and the AR0387 group contained four biological replicates. A total of 40 mice were used.

### Generation of mutant strains of genes B9J08_001458 (*SCF1*) and B9J08_004112 (*ALS5*)

#### Plasmid construction

The plasmids used in this study were propagated in *E. coli* TOP10F’chemically competent cells. Bacterial transformations were carried out by heat shock at 42°C for 45 sec using 30 μl of competent cells, and subsequent cooling on ice for 2 min. The transformants were selected on solid LB (Sigma, Fisher Scientific) medium (agar 15%, Bacto™ Agar, BD) supplemented with ampicillin (100 μg/ml). To construct the deletion mutants, we utilized the SAT1 flipper tool^52^. Specifically, the upstream and downstream regions of the genes of interest were amplified from the genomic DNA of *C. auris* strain B8441 with primers listed in Table S1, and cloned into pSFS2 in a homodirectional way so that they flanked the nourseothricin resistance marker (*SAT1*) and the *FLP* recombinase gene. To generate the B9J08_001458 deletion cassette, the upstream homologous region was cloned into the XhoI/KpnI-HF (NEB) digested pSFS2 plasmid using NEBuilder HiFi (NEB) as per manufacturer instructions. The resulting constructs were isolated from the transformants, digested with SacI-HF and NotI-HF and the downstream region was cloned into the digested plasmid. To generate the B9J08_004112 deletion cassette, the upstream homologous region was cloned into the ApaI/KpnI-HF (NEB) digested pSFS2 plasmid. Resulting constructs were isolated from the transformants, digested with SacII and NotI-HF and the downstream region was cloned into the digested plasmid. All inserts of the plasmids were verified by sequencing (Mix2Seq, Eurofins genomics). To produce a linear deletion cassette, each plasmid was digested with KpnI-HF and SacII for the B9J08_004112 deletion cassette and with StuI and ScaI for the B9J08_001458 deletion cassette. Primers used to verify correct insertion of the upstream and downstream regions are listed in Table S1.

#### Strain construction

For strain construction, *C. auris* cells were prepared as described by Carolus *et al.*^53^. For electroporation, 40 μl of competent cells was mixed with the transformation mixture and transferred to 2 mm electroporation cuvettes (Pulsestar, Westburg). The transformation mixtures comprised 3 μl of 4 μM Alt-R™ S.p. Cas9 Nuclease V3, 3.6 μl of duplexed gene-specific Alt-R® CRISPR-Cas9 crRNA (IDT) with Alt-R® CRISPR-Cas9 tracrRNA (IDT) and 500 ng of the linearized constructed pSFS2 variant for each gene as donor DNA. A single pulse was given at 1.8 kV, 200 W, 25 mF, and the transformation mixture was immediately transferred to 2 ml YPD in test tubes and incubated for 4 h at 37°C at 150 rpm. Cells were collected by centrifugation for 5 min at 5000 g, resuspended in 100 μl YPD and plated on YPD agar containing 200 μg/ml of nourseothricin (Jena Bioscience). The sequences of the crRNA are listed in Table S1. Correct transformants were screened by colony PCRs, using the Taq DNA Polymerase (NEB) and primers that bind in the deletion cassette and outside of the homologous regions upstream and downstream (Table S1). Null mutants of B9J08_001458 (*SCF1*) and B9J08_004112 (*ALS5*) were generated in triplicate (3 independent transformants; Δ1-Δ3) in both wild-type backgrounds (AR0387 and AR0382) and evaluated for biofilm formation but only one representative mutant was randomly selected for subsequent analysis (Fig. S2).

#### Evaluation of potential complementary roles for the Scf1 and Als5 adhesins in surface adhesion and biofilm formation

To determine the impact of gene deletions on adherence and biofilm formation and whether there are adherence complementary roles for the Scf1 and Als5 adhesins, mutant strains were compared to the wild-type strain individually and in combination in biofilm assays based on assessment of metabolic activity. Biofilms were grown by seeding 200 µl of 1x10^6^ cells/ml cell suspensions of each strain in flat-bottom 96-well polystyrene microtiter plates; for combination biofilms, mixed solutions of 100 µl of 1x10^6^ cells/ml cell suspensions of each mutant were used. Following incubation at 37°C for 24 h, wells were washed with PBS and biofilms evaluated using the MTS metabolic assay (Promega) as per manufacturer recommendation. Color intensity was measured at 490nm using a Cell Imaging Multi-Mode Reader (Cytation 5, Biotek). Assays were performed on 3 separate occasions, each using 4 technical replicates.

### Evaluation of potential complementary roles for the Scf1 and Als5 adhesins in cell-cell adherence and coaggregation

The contribution of adhesins to cell-cell interaction and coaggregation was comparatively assessed based on formation of cell aggregates. For coaggregation assays, cell suspensions of wild-type strain and mutant strains were suspended in PBS to final density of 5x10^8^ cells/ml in 5 ml plastic tubes, and suspensions vigorously vortexed for 1 min. To evaluate adherence complementation of adhesins, suspensions of both mutants at 2.5x10^8^ cells/ml were equally mixed and vortexed. Tubes were placed upright at RT and cell aggregation was monitored and imaged. Additionally, sedimentation rate of formed cell aggregates was also measured based on drop in absorbance readings of cell suspensions. For these experiments, aliquots from cell suspensions were measured at 600nm every 10 min for up to 2 h in a BioTek 800 TS absorbance reader. Sedimentation rate was calculated as the percent reduction in absorbance at each timepoint compared to the initial reading.

#### Confocal laser scanning microscopy (CLSM) and scanning electron microscopy (SEM) analysis of biofilms of wild-type and the Δ*scf1* and Δ*als5* mutants grown individually and in combination

For CLSM, biofilms of wild-type and mutant strains individually and in combination were grown on glass coverslip-bottom dishes (MatTek Co., Ashland, MA) for 24 h; biofilms were rinsed in PBS then stained with a concanavalin-A conjugated to Alexa 647 (Invitrogen) (50 µg/ml) for 45 min at 37°C. Biofilms were visualized using an inverted confocal laser scanning microscope (T2i, Nikon) and images analyzed using Imaris 9.3 Arena software and ImageJ. For SEM, biofilms were grown on coverslips for 24 h at 37°C then fixed in 2% paraformaldehyde-2.5% glutaraldehyde, post-fixed with 1% osmium tetroxide, serially dehydrated in ethyl alcohol (30-100%) and critical-point dried. Samples were carbon-coated and observed with Quanta 200 SEM (FEI Co.) and images processed using Adobe Photoshop software.

### *In vivo* evaluation of AR0382 and AR0387 wild-type strains and Δ*scf1* mutant in a mouse subcutaneous catheter model

Based on observed *in vitro* biofilm deficiency of the Δ*scf1* mutant, its ability to form biofilm on catheters *in vivo* was evaluated in the subcutaneous catheter model. The adherence of Δ*scf1* was compared to both the aggregative (AR0382) and non-aggregative (AR0387) wild-type strains. For these experiments, catheter fragments inoculated with the strains *in vitro* were implanted in animals as described above. Biofilms were allowed to form within catheters for 72 h then animals were euthanized and catheters harvested. To visualize the biofilms formed within the catheter lumen, catheters from each group were cut longitudinally to expose the lumen, fixed and processed for SEM analysis as described above. Catheters from 6 mice were analyzed and representative images presented.

### Comparative evaluation of cell-cell adhesion forces between cells of *C. auris* strains using single-cell force spectroscopy (SCFS)

SCFS was employed to measure single cell-cell adhesion forces among cells of wild-type and the two mutants^54–56^. For these studies, a single live cell was attached to a polydopamine-coated tipless AFM cantilever and approached toward another single cell, previously immobilized on a dish. The retraction and approach movement of the cell probe was monitored and force-distance curves recorded, allowing quantification of the forces driving intercellular adhesion. Triangular tipless cantilevers (NP O10, Bruker) were immersed for 1 h in Tris-buffered saline solution (50 mM Tris, 150 mM NaCl, pH 8.5) containing 4 mg/ml of dopamine hydrochloride, rinsed with Tris-buffered saline solution and mounted on the AFM setup for cell probe preparation. Calibration of the probe was performed prior to the AFM experiment and its nominal spring constant determined by the thermal noise method. *C. auris* cells were grown overnight in liquid YPD at 37°C, 150 rpm, harvested by centrifugation, washed three times in 1X PBS and finally diluted 1000-fold. Cell suspensions were allowed to adhere to polystyrene dishes for 20 min and dishes washed three times then filled with 2 ml of 1X PBS before being transferred to the AFM setup. SCFS measurements were performed at RT in 1xPBS, using a Nanowizard 4 AFM (JPK Instrument, Berlin, Germany). The cell probe was prepared by bringing the polydopamine-coated cantilever into contact with an isolated cell and, once the probe was retracted, its attachment to the cantilever was confirmed using an inverted optical microscope. The cell probe was then positioned over an immobilized cell and force maps of 16x16 pixels were recorded on top of it, using a contact force setpoint of 250 pN, a constant approach and retraction speeds of 1 µm/s and an additional pause at contact of 1 s. Adhesion forces were extracted from force-distance retraction curves by considering the rupture event for which the adhesion force was maximal, for every curve.

### Data analysis

Statistical analysis of biofilm growth was performed using R statistical programming software. Statistical analysis of SCFS data was performed with Origin software (OriginPro 2021). To compare differences among strains in *in vitro* biofilm forming capabilities and cell-cell adhesion force and frequency, a one-way ANOVA with Tukey’s post host test was used. *P* values less than 0.05 were considered significant. Two-sample t-tests were used to compare absorbance values, adhesion force and adhesion frequency between AR0382 and AR0387 strains. Ggplot2 and ggpubr packages were used to construct models for figure construction.

## DATA AVAILABILITY

Upon acceptance and prior to publication, all of the raw sequencing reads from this study will be available at the NCBI Sequence Read Archive (SRA). All strains generated in this study will be made available upon request from authors.

## ACKNOWLEDGEMENTS

The work in this publication was supported by the National Institute of Allergy and Infectious Diseases of the NIH under award number R01AI130170 (NIAID) to M.A.J-R, and NIH grant U19 AI110820 to V.M.B. This work was also supported by the University of Maryland Baltimore, Institute for Clinical & Translational Research (ICTR), the Fund for Scientific Research, Flanders, research community on biofilms (FWO #W000921N), and the Belgian National Fund for Scientific Research (FNRS). We would like to thank Miles Delmar and the UMB Electron Microscopy Core for SEM imaging.

## Author Contributions

M.A.J-R. and P.V.D. conceived and designed this research, M.A.J-R., V.M.B. and P.V.D. provided funding; T.W.W., D.M-J., D.S., H.C., C.M., A.A. and T.O.P. performed experiments; M.A.J-R., T.W.W., D.S., T.O.P., V.M.B., P.V.D., D.M-J. and Y.F.D. analyzed data; M.A.J-R., T.W.W., D.S., T.O.P., H.C. and V.M.B. wrote the paper; M.A.J-R. oversaw the entire study. All authors read and approved the manuscript

## Supplemental Material

**Supplemental Figure S1. Probing *C. auris* cell-cell adhesion using single-cell force spectroscopy.** AFM setup used for single-cell force spectroscopy experiments. A single live *C. auris* cell was attached to a tipless AFM cantilever previously functionalized with polydopamine. This cell probe was moved toward another single *C. auris* cell immobilized on a polystyrene dish and force-distance curves were recorded, allowing quantification of the intercellular adhesion forces.

**Supplemental Figure S2. Evaluation of biofilm formation by the 3 mutant strains generated for the *ALS5* and *SCF1* genes *(Δ*1-*Δ*3).** A measurement of the metabolic activity of 24 h biofilms based on values of OD_490_ comparing all generated mutant strains to the wild-type. Boxplots show mean and SEM of *n* = 3 biological replicates, each as an average of 4 technical replicates. Statistical analysis was performed by one-way ANOVA and post-hoc Tukey test with *p*-values representing significant differences. *P*=1.64×10^-3^, 1.55×10^-3^, 2.18×10^-3^, 2.02×10^-4^, 3.70×10^-4^, 3.50×10^-4^, 4.83×10^-4^, 5.17×10^-5^, 3.35×10^-4^, 3.17×10^-4^, 4.37×10^-4^, 4.71×10^-5^ **0.001 < *P* ≤ 0.01, \*\*\**P* < 0.001.

**Supplemental Figure S3. Comparative evaluation of biofilm formation, aggregation and cell-cell adhesion force by the wild-type AR0382 (aggregative) and AR0387 (non-aggregative) phenotypes. (A)** Metabolic activity of 24 h biofilms based on measurements of OD_490_, optical density. Values are means plus standard errors of the means (error bars). Statistical analysis was performed by an unpaired two-sided t-test. Bar-graphs shows mean and SEM of *n* = 3 biological replicates, each as an average of 4 technical replicates. *P* = 2.243×10^-5^. **(B)** Aggregation assays, following vigorous vortexing of cell suspensions comparing cell aggregates of AR0382 and AR0387. Bright-field microscopy (lower panel) of aliquots of cell suspensions demonstrating presence of aggregates of AR0382 cells compared to singly suspended cells of AR0387. **(C**) Measurement of rate of cell sedimentation by absorbance readings of OD_600_ of wild-type strains AR0382 and AR0387 over 2 h following vigorous vortexing. Values represent mean OD and SEM of three technical replicates. **(D)** Single-cell force spectroscopy of *C. auris* cell-cell adhesion. Adhesion force histograms with representative retraction profiles (inset) obtained for the interaction between AR0382 wild-type cells and the interaction between AR0387 cells; 2 representative cell pairs are shown for each strain. **(E)** Adhesion force boxplots depicting *n* = 6 and *n* = 4 cell pairs for AR0382 and AR0387 respectively. Statistical analysis was performed by an unpaired two-sided t-test. *P* = 4.21×10^-2^ **(F)** As in **(E)**, adhesion frequency boxplots show interactions between *n* = 7 cell pairs for both strains. *P* = 8.06×10^-6^. Red stars represent the mean values, red lines are the medians, boxes are the 25−75% quartiles and whiskers the standard deviation from mean. *0.01 < *P* ≤ 0.05, \*\*\**P* < 0.001.

**Table S1.** Primers used in this study

**Table S2.** Differentially expressed genes between AR0382/AR0387 during *in vitro* biofilm growth (FDR <0.01, LFC >= |1.0|)

**Table S3.** Differentially expressed genes between AR0382/AR0387 during *in vivo* biofilm growth (FDR <0.01, LFC >= |1.0|)

**Table S4.** List of genes that are more highly expressed in AR382 under both *in vitro* and *in vivo* biofilm conditions (FDR <0.01, LFC >= |1.0|)

## Notes

### Competing Interest Statement

The authors have declared no competing interest.

