## Supplemental Material for "Functional Redundancy in *Candida auris* Cell Surface Adhesins Crucial for Cell-Cell Interaction and Aggregation"

### Supplementary Table 4 (S4). Primers used in this study.

| Marker | Purpose | Primer name | Sequence (5'-3') |
|---|---|---|---|
| *B9J08_004112* | US_cloning | CauALS5_US_GIB_F | AACTTCCTCGAGGGGGGGCCGAAAGATGATGGGAAACAAGGTGAAG |
| | US_cloning | CauALS5_US_GIB_R | AGGGAACAAAAGCTGGGTACCTCACCACGAGACGGGAGC |
| | DS_cloning | CauALS5_DS_GIB_F | GCGAATTGGAGCTCCACCGCGGTCCCCAGGTGCTATTTCTTG |
| | DS_cloning | CauALS5_DS_GIB_R | AGATCCACTAGTTCTAGAGCACGTGAGCTTTTATGATACCTAC |
| | crRNA | ALS5_crRNA | GTACTCAGGCGAAAACATCG |
| | | ALS5US_chF | GTCATTCTTCCTTGTTTCCG |
| | | ALS5DS_chR | CATTTGCAGAGAAGAAATCTGCGC |
| *B9J08_001458* | US_cloning | Gbson_RBT1US_F | TAGAAAGTATAGGAACTTCCTGTGGAGGTGAAGTTTTAAGATAGAG |
| | US_cloning | Gbson_RBT1US_R | AGGGAACAAAAGCTGGGTACGCTCGCCGCTCACAATG |
| | DS_cloning | Gbson_RBT1DS_F | CTATAGGGCGAATTGGAGCTGTCGGGATTGTGGGAATTC |
| | DS_cloning | Gbson_RBT1DS_R | AGATCCACTAGTTCTAGAGCTTCTAATGACTGATACTCATACTTTC |
| | Verify deletion | RBT1US_chF | ATGTGCTTCTTCTGGGTCTTTTG |
| | Verify deletion | RBT1DS_chR | GCGATAGGAGACGATGTTGATAAC |
| | crRNA | RBT1_crRNA | CTAGGTCCACTAGGTCCACT |
| pSFS2 | PCR/Seq/verify deletion | pSAT1_US-cPCR_F | CTAACGATGCATACGACTACATC |
| | PCR/Seq/verify deletion | pSAT1_DS-cPCR_R | ACATATGTGAAGTGTGAAGGGGG |

**Supplemental Figure S1. Comparative evaluation of biofilm formation, aggregation and cell-cell adhesion force by the wild-type AR0382 (aggregative) and AR0387 (non-aggregative) phenotypes.** **(A)** Metabolic activity of 24 h biofilms based on measurements of $OD_{490}$, optical density. Values are means plus standard errors of the means (error bars). Statistical analysis was performed by an unpaired two-sided t-test. Bar-graphs shows mean and SEM of $n$ = 3 biological replicates, each as an average of 4 technical replicates. $P = 2.243\times10^{-5}$. **(B)** Aggregation assays, following vigorous vortexing of cell suspensions comparing cell aggregates of AR0382 and AR0387. Bright-field microscopy (lower panel) of aliquots of cell suspensions demonstrating presence of aggregates of AR0382 cells compared to singly suspended cells of AR0387. **(C)** Measurement of rate of cell sedimentation by absorbance readings of $OD_{600}$ of wild-type strains AR0382 and AR0387 over 2 h following vigorous vortexing. Values represent mean OD and SEM of three technical replicates. **(D)** Single-cell force spectroscopy of *C. auris* cell-cell adhesion. Adhesion force histograms with representative retraction profiles (inset) obtained for the interaction between AR0382 wild-type cells and the interaction between AR0387 cells*;* 2 representative cell pairs are shown for each strain. **(E)** Adhesion force boxplots depicting $n$ = 6 and $n$ = 4 cell pairs for AR0382 and AR0387 respectively. Statistical analysis was performed by an unpaired two-sided t-test. $P = 4.21\times10^{-2}$ **(F)** As in **(E)**, adhesion frequency boxplots show interactions between $n$ = 7 cell pairs for both strains. $P = 8.06\times10^{-6}$. Red stars represent the mean values, red lines are the medians, boxes are the 25−75% quartiles and whiskers the standard deviation from mean. $^*0.01 < P \leq 0.05$, $^{***}P < 0.001$.

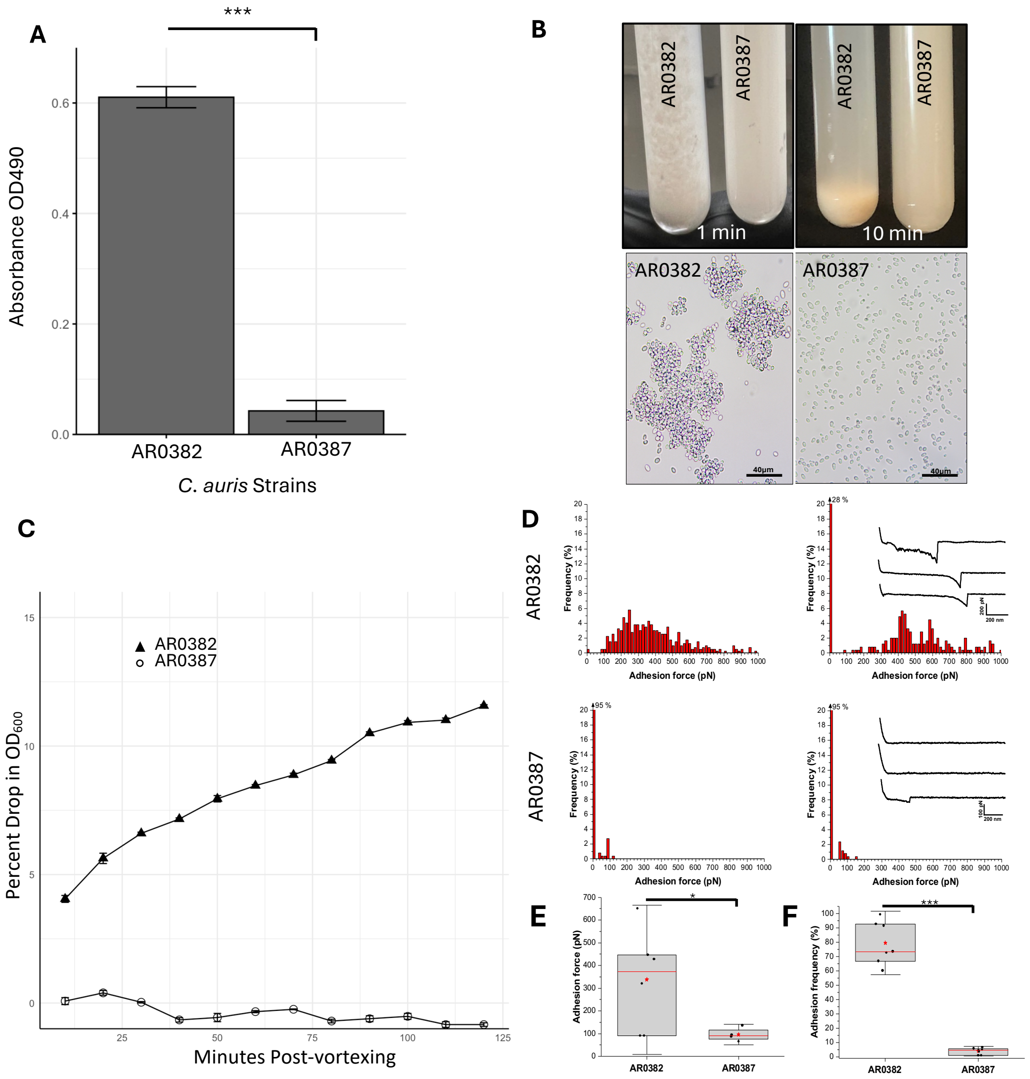

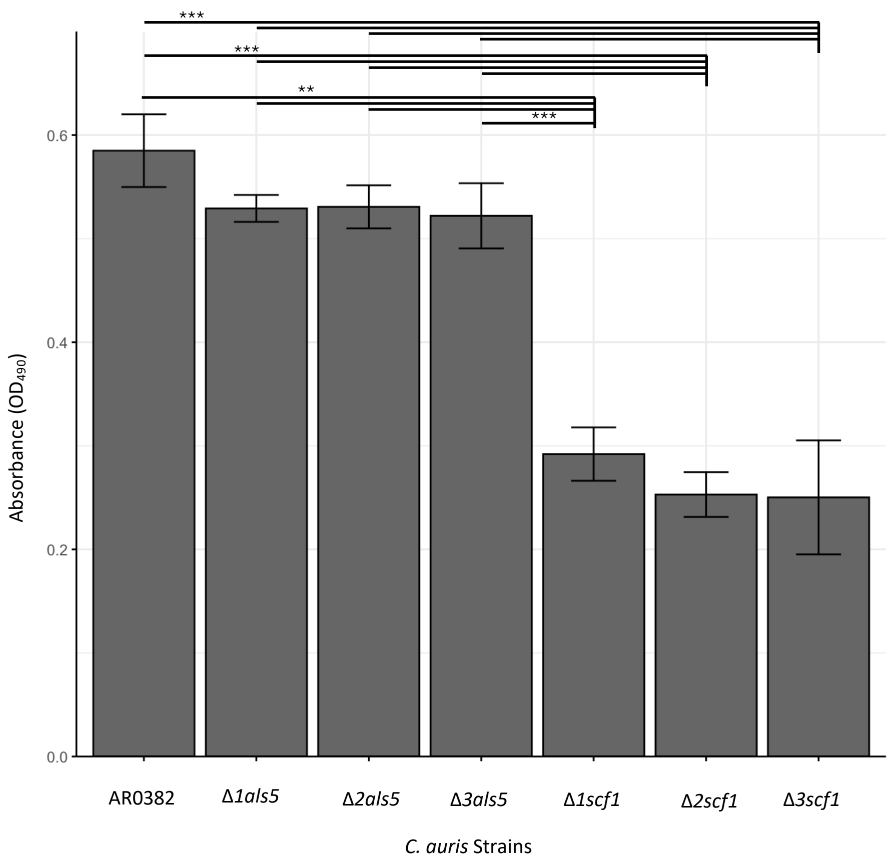


**Supplemental Figure S2. Evaluation of biofilm formation by the 3 mutant strains generated for the *ALS5* and *SCF1* genes (*Δ*1-*Δ*3).** A measurement of the metabolic activity of 24 h biofilms based on values of $OD_{490}$ comparing all generated mutant strains to the wild-type. Boxplots show mean and SEM of *n* = 3 biological replicates, each as an average of 4 technical replicates. Statistical analysis was performed by one-way ANOVA and post-hoc Tukey test with *p*-values representing significant differences. $P=1.64\times10^{-3}$, $1.55\times10^{-3}$, $2.18\times10^{-3}$, $2.02\times10^{-4}$, $3.70\times10^{-4}$, $3.50\times10^{-4}$, $4.83\times10^{-4}$, $5.17\times10^{-5}$, $3.35\times10^{-4}$, $3.17\times10^{-4}$, $4.37\times10^{-4}$, $4.71\times10^{-5}$ **$0.001 < P \leq 0.01$, ***$P < 0.001$.
